# The whole genome sequence and mRNA transcriptome of the tropical cyclopoid copepod *Apocyclops royi*

**DOI:** 10.1101/502997

**Authors:** Tue Sparholt Jørgensen, Bolette Lykke Holm Nielsen, Bent Petersen, Patrick Denis Browne, Benni Winding Hansen, Lars Hestbjerg Hansen

## Abstract

Copepoda is one of the most ecologically important animal groups on Earth, yet very few genetic resources are available for this Subclass. Here, we present the first whole genome sequence (WGS, acc. UYDY01) and the first mRNA transcriptome assembly (TSA, Acc. GHAJ01) for the tropical cyclopoid copepod species *Apocyclops royi*. Until now, only the 18S small subunit of ribosomal RNA gene and the COI gene has been available from *A. royi*, and only one other cyclopoid copepod had WGS resources available. Overall, the provided resources are the 7^th^ copepod species to have WGS available and the 19^th^ copepod species with TSA information available. We analyze the length and GC content of the provided WGS scaffolds as well as the coverage and gene content of both the WGS and the TSA assembly. Finally, we place the resources within the copepod order Cyclopoida as a member of the *Apocyclops* genus. We estimate the total genome size of *A. royi* to 450 Mb, with 181 Mb assembled nonrepetitive, 76 Mb assembled repeats and 193Mb unassembled sequence. The TSA assembly consists of 29,737 genes and an additional 45,756 isoforms. In the WGS and TSA assemblies, >80% and >95% of core genes can be found, though many in fragmented versions. The provided resources will allow researchers to conduct physiological experiments on *A. royi*, and also increase the possibilities for copepod gene set analysis, as it adds substantially to the copepod datasets available.

## Introduction

Copepods are among the most numerous animals on Earth, and the ecology, behavior, biotechnological and aquaculture potential of copepods has been scrutinized for decades. Yet very few molecular resources are available for the subclass Copepoda. *Apocyclops royi* is an omnivorous cyclopoid copepod found in estuaries, brackish-water aquaculture ponds and in freshwater areas in tropical regions [1–4]. *A. royi* is a relatively small egg-carrying copepod with a prosome length of 0.5 mm (Fig 1A) [1, 2]. It has a life cycle of 7-8 days [5], and can tolerate temperatures of 15-35 °C [6], and salinities of 0-35 psu [7]. In a recent publication, we report its ability to biosynthesize the polyunsaturated fatty acid Docosahexaenoic acid (DHA) from *alpha*-Linolenic acid [8, 9] which makes *Apocyclops royi* an interesting organism for copepod physiological studies.

**Fig. 1.**
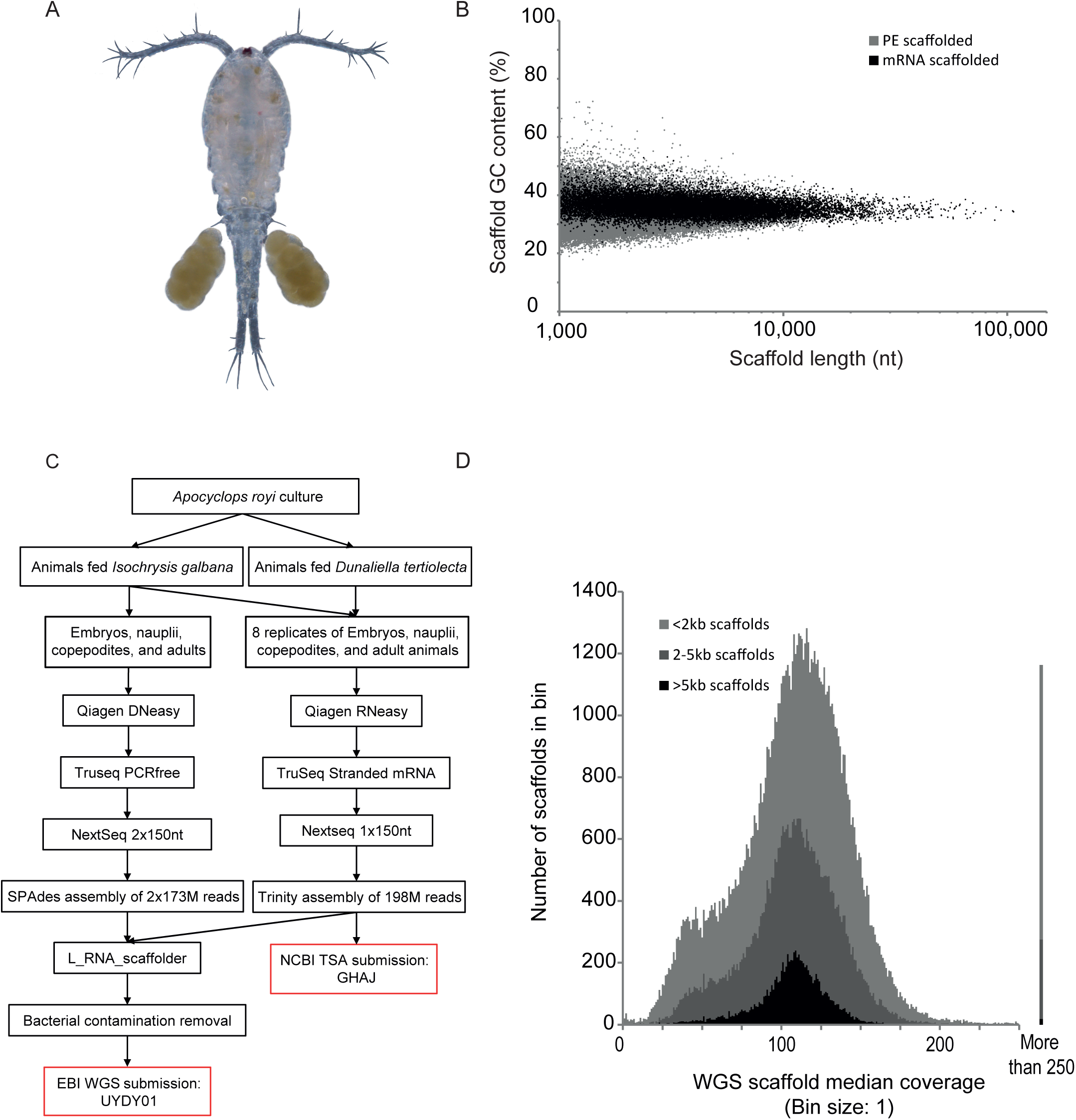
A, composite picture of a female *Apocyclops royi* with egg sacks from the culture used for experiments. B, WGS assembly GC% plotted against scaffold length for mRNA scaffolded sequences (black) and not mRNA scaffolded sequences (grey). Each dot represents one scaffold. C, workflow used in the present study from culture to data deposition. D, median coverage estimation for the WGS assembly in the three scaffold length subsets <2kb, 2-5kb, and >5kb. For each, a maximum number of scaffolds are seen in bins with coverage ca 110. Less than 1% of scaffolds have a coverage higher than 250 (illustrated on the right hand side of the plot). We chose to use median values to minimize the impact of highly covered regions, which are regularly seen in WGS datasets and which are likely owing to repetitive sequence.

Copepod genomes are infamously difficult to assemble [10, 11]. This is likely caused by high repetitiveness, a low GC content of around 30% [12] and very variable genome sizes [13], which means that it is difficult to assess the costs before undertaking whole genome sequencing (WGS). This is compounded by the often small physical size of the animals, which makes it necessary to use a collection of animals rather than a single individual for nucleic acid purification, adding to the complexity of genome assembly. Modern genome assembly pipelines and data generation workflows are optimized for mammalian genome assembly, and any deviation from mammalian like genomes are likely to result in lower quality assembly. Crucially, the total genome size often differs substantially from the assembly size, as repetitive DNA is collapsed or remains unassembled. Transcriptome assemblies, however, are significantly easier to obtain, as many of the clade-specific limitations of copepod WGS are overcome by focusing on mRNA. Here, the highly repetitive regions are not transcribed or are removed post-transcriptionally and the assembly process is simpler as the remaining repetitive regions are dealt with simplistically [14]. A recent paper presents a good example of a high quality transcriptome from a copepod where WGS information was not available [15]. A lot of information is however not captured by a transcriptome. For example, intron sizes and repeat structure can be derived from a genome assembly, but not from a transcriptome, which also fails to capture genes not constitutively expressed or which are expressed only in few cells or in certain tissues.

For evolutionary analysis relying on existing DNA databases, it is imperative to have a diverse range of genomic information available. As of now, only one cyclopoid copepod genome is available, namely the high quality WGS assembly of *Oithona nana* [16]. Further, only seven copepod species have available WGS information, and only 19 copepod species have available TSA information, including the *A. royi* datasets. With the presented *A. royi* genome and transcriptome, we expand the possibilities for studies centered on *A. royi* physiology and improve the possibility for large scale phylogenetic and evolutionary studies. Further, our high-quality short read resources may prove pivotal in error correcting future genome projects which will utilize error prone third generation DNA and RNA sequencing.

## Methods and Materials

### Organism origin and derivation

An overview of the experimental and bioinformatical workflow can be seen in Fig 1C. Animal husbandry, sampling, RNA purification, RNA sequencing strategy and initial RNA data processing is also described in a recent paper which used the mRNA dataset presented here to analyze the fatty acid metabolism genes and differential expression based on feeding regime [9] (in review). Briefly, an *Apocyclops royi* animal culture obtained from Tungkang Biotechnology Research Center in Taiwan was split in two which were kept in 100 L tanks on separate microalga feeding regimes. One was fed *Isochrysis galbana* and the other was fed *Dunaliella tertiolecta*.

#### Sampling

Animals from each feeding regime were sampled as described in [9]. Briefly, animals were starved for 2 h to empty their guts and a 53 µm filter was used to separate all the life stages (nauplii, copepodites and adults) from the sea water. Four analytical replicates were made for each feeding regime, each consisting of hundreds to thousands of individuals. Animals were flushed with 0.2um filtered seawater (32ppt) until visual inspection showed very little particular contaminating matter. The remaining seawater was aspirated with a homemade small tip pasteur pipettor to ensure that animals were not removed during this step. A volume of 200 µl of RNAlater was added to the replicates of animals fed *I. galbana* and 500 µl of RNAlater was added to the replicates of animals fed *D. tertiolecta* to ensure a factor of at least 1:10 of animals in RNAlater. Samples were kept in a fridge for 24 H and frozen until use. These samples were used for both RNA and DNA extractions.

### Sequencing methods and preparation details

#### Nucleic acid extraction

as described in [9], RNA was extracted with RNeasy (Qiagen, Venlo, Nederlands) according to protocol. Before extraction, all RNAlater was removed and the animal tissue was disrupted with a 1.5 mL RNase-Free Pellet Pestle (Kimble Chase, Vineland, New Jersey, USA) mounted on a Kontes Pellet Pestle motor (Kimble Chase, Vineland, New Jersey, USA) for 1 min on ice in 20 µl buffer RTL, before adding the remaining 320 µl Buffer RTL.

DNA was extracted from replicate 2 of animals fed *I. galbana* using the DNeasy blood and tissue kit from Qiagen according to protocol. Briefly, tissue from thousands of animals was disrupted manually with a 1.5 ml RNase-Free Pellet Pestle (Kimble Chase, Vineland, New Jersey, USA) to prevent unnecessary DNA shearing. The ground tissue was incubated for four hours at 56 °C in lysis buffer with Proteinase K according to protocol, vortexing every 15-30 min. A Qubit 3.0 fluorometer (Thermo Fisher Scientific, Waltham , MA, USA) was used to determine the DNA and RNA concentrations.

#### Sequencing library construction

The RNA sequencing library strategy is described in [9]. Briefly, an mRNA sequencing library was produced for each of the eight replicates with the Truseq stranded mRNA kit (Illumina, San Diego, California, United States) and SuperscriptII reverse transcriptase (Thermo Fisher Scientific, Waltham, MA, USA). 1 µg total RNA was used for each of the eight mRNA library preparations. DNase was not used to avoid breakdown of long transcripts and because the stranded protocol minimizes the influence of DNA contamination. The efficiency of the protocol was assessed using the directionality of reads. A PCR-free DNA sequencing library was produced using the Illumina TruSeq PCR-Free kit (Illumina, San Diego, California, United States) according to protocol. DNA was sheared in a Covaris E210 with the following settings: Intensity: 4, Duty cycle 10%, Cycles per burst: 200, Treatment time: 70 s intended to produce fragments of 350nt.

The library cluster forming molarity of all samples was evaluated using the KAPA qPCR system (Roche, Basel, Switzerland) and samples were run on a Bioanalyzer 2100 (Agilent Technologies, Santa Clara, CA, USA) to evaluate the fragment length.

#### Sequencing

The eight mRNA libraries were pooled equimolarly and run on a single Illumina Nextseq 1×150nt mid output flowcell as described in [9]. The PCR-free DNA library was run on a single Illumina Nextseq 2×150nt mid output flowcell.

#### Data processing methods

Initial data handling and basic statistics were carried out using Biopieces (Hansen, MA, www.biopieces.org, unpublished). Raw illumina reads were trimmed using Adapterremoval v. 2.0 [17] with the following parameters for the RNA data “--trimns --trimqualities” and standard parameters for paired end data for the PCR-free WGS reads. Trinity v. 2.5.1 [14] was used to assemble pooled mRNA reads from all eight sequenced replicates with the following parameters: “--SS_lib_type R --trimmomatic --single”. Transcripts shorter than 500nt were discarded and PhiX contigs removed by BLAST [18] in CLCgenomics 11.0 (Qiagen, Venlo, Nederlands). The PreQC system from the SGA pipeline was used to estimate the total genome size based on read k-mer spectra [19]. SPAdes v. 3.11 [20] with the auto-selected k-mer sizes 21, 33, 55, and 77 was used to assemble the PCR-free WGS reads on the Computerome supercomputer on a 1TB RAM node. The SPAdes log can be found in Supplementary Material 1. The WGS assembly was scaffolded using the mRNA TSA assembly and the L_RNA_scaffolder program [21].

#### Contamination removal

Because whole animals were used for the WGS data generation, it is expected that bacterial symbionts also contributed DNA to the sequencing libraries. In order to remove any sequence of bacterial origin from the genome assembly, we first masked all scaffolds using Repeatmodeler and Repeatmasker (v. 4.0.7) [22, 23]. Repeats from RepeatModeler and the Arthropoda and ancestral (shared) repeats from repbase v. 22.05 (downloaded 2017-06-02) were used to mask scaffolds. The masked scaffolds were searched against the RefSeq database of representative prokaryotes (downloaded 2017-03-23) using the build-in BLAST in CLCgenomics 11.0. Scaffolds with BLAST hits longer than 500nt without mRNA proof were removed from the assembly. The output from a second round of Repeatmasker run on the assembly without contamination was used to estimate the assembled repetitive and non-repetitive fractions of the WGS assembly.

The sequencing depth was estimated by mapping all reads on assemblies using Bowtie2 [24] (v. 2.3.4, switches: --local --no-unal) and extracting the median coverage of each transcript (TSA assembly) or scaffold (WGS assembly) using Samtools [25] (samtools view |samtools sort -|samtools depth -aa -) and a custom pyton script which can be found in supplementary material 2. Both the mRNA TSA assembly and the WGS genome assembly were evaluated using the BUSCO Universal Single-Copy Ortholog v.2 [26]. In order to obtain a 18S rRNA gene sequence, paired reads from the WGS dataset was mapped on the partial cyclopoida genes from the PopSet 442571920 [27] using Bowtie2. The read pairs where at least one read mapped were then extracted and assembled using SPAdes v. 3.13 and the resulting 18S rRNA gene sequence was aligned to the reference sequences and trimmed using CLCgenomics 11.0. A neighbor-joining phylogram was constructed in CLC genomics 11.0 using 1000 bootstraps.

### Results and discussion

After quality and adapter trimming, the sequencing yielded 173,365,491 PCR-free WGS read clusters (346,730,982 reads) and 203,548,224 mRNA derived reads constituting 52 Gbases and 31 Gbases of data, respectively. In total, 99.9% and 97.1% of reads was left after quality and adapter trimming and filtering, respectively.

The TSA assembly yielded 100.7 Mb in 29,730 genes and additionally 45,747 alternative isoforms giving a total of 75,477 transcripts. The WGS assembly yielded 143,521 contigs in 97,072 scaffolds comprising a total length of 257.5 Mb, while 83.6% of sequencing reads mapped back to the assembly (data not shown). The size of the assembly is similar to other copepod WGS datasets, but three times larger than the *Oithona nana* assembly, which is the only other cyclopoid copepod WGS assembly available.

After bacterial contamination removal, the WGS assembly consists of scaffolds up to 116 Kb in length, with an average GC% of 33.5% (Fig. 1B), which is similar to other available copepod WGS assemblies, such as *Acartia tonsa* [13]. The uniformity of the length and GC% in Fig. 1B suggests that most contaminants are not present in the assembly, as bacteria and other contaminants would likely have a different pattern of distribution of scaffold length and GC%. For example, we removed several contigs in the size range 100 kb to 1 Mb, all with a GC content between 56 % and 58 % and highly similar to known bacterial sequences. In order to estimate the genome size of *A. royi* including the unassembled and repetitive fraction, we used the preQC program which has previously been used for copepod genome size estimation [13]. The result shows that the expected complete genome size of *A. royi* is 450 Mb (supplementary material 3 preQC report). Of this, 181 Mb are assembled nonrepetitive sequence, 76 Mb are assembled repeats and 193Mb are unassembled sequence (Repeatmodeler output can be found in Supplementary material 4). Much of the unassembled sequence is can presumably be found in scaffolds smaller than the 1kb cutoff, though repeats also would be collapsed in these scaffolds. In a recent publication on the *Acartia tonsa* WGS assembly, the genome sizes of all copepod WGS projects was estimated and in all cases showed that less than half of the expected genome size was included in the WGS assembly [13]. The difference between the assembled and the actual size of the *A. royi* genome is thus expected, similar to the differences in other species, and hypothesized to be largely caused by unassembled repetitive/non-coding regions or collapsed scaffolds [13]. For example, if a repeat of 500 nt is found 1.000 times scattered throughout the genome, the sequence is unlikely to show up more than once in the assembly, which means that the assembly size is 500.000 nt smaller than the template genome. This repeat scaffold would then have 1.000 times higher coverage than the non-repetitive fraction of the genome assembly.

In figure 1D, a histogram of the median scaffold coverage (binsize 1) between 1 and 250 show that the largest amount of scaffolds in each of the three scaffold length fractions have a coverage of ca 110 (Supplementary Material 5). This result fits the simplistic coverage estimation: 52Gb of reads should give a coverage of ca 115 on a 450Mb genome. We chose to use median rather than mean values to minimize the impact of scaffold regions with extremely high coverage, which are often seen in copepod assemblies and potentially are the result of assembled repetitive sequence. In the smaller scaffold size fractions <2 kb and 2-5 kb, a distinct shoulder is observed at coverage ca 35. In Fig. 1D, scaffold bins with a coverage between 0 and 250 are shown, but many scaffolds had a higher coverage than 250. These were collected in a separate bin (>250) which is displayed on the right hand side of Fig 1D, and likely constitute many of the repeated regions in the genome. In total, only 1.2% of scaffolds have a coverage higher than 250. It is generally recommended to produce WGS assemblies from datasets with coverage of ca. 100, which the results in Fig. 1D confirm was achieved. By mapping the mRNA derived reads to the transcripts of the TSA dataset, we similarly produced an overview of the median coverage of transcripts (Fig 2A). Importantly, the coverage in transcriptomes are not similar to those in WGS assemblies in that differential expression of genes means that a uniform coverage is not expected. As a result of this, the range of transcript median coverage bins seen in Fig. 2A had to accommodate a median coverage distribution from near-zero to more than 4,000,000 though >99% of transcripts had a median coverage of less than 1000 (Suplementary Material 5 mapping table).

**Fig. 2.**
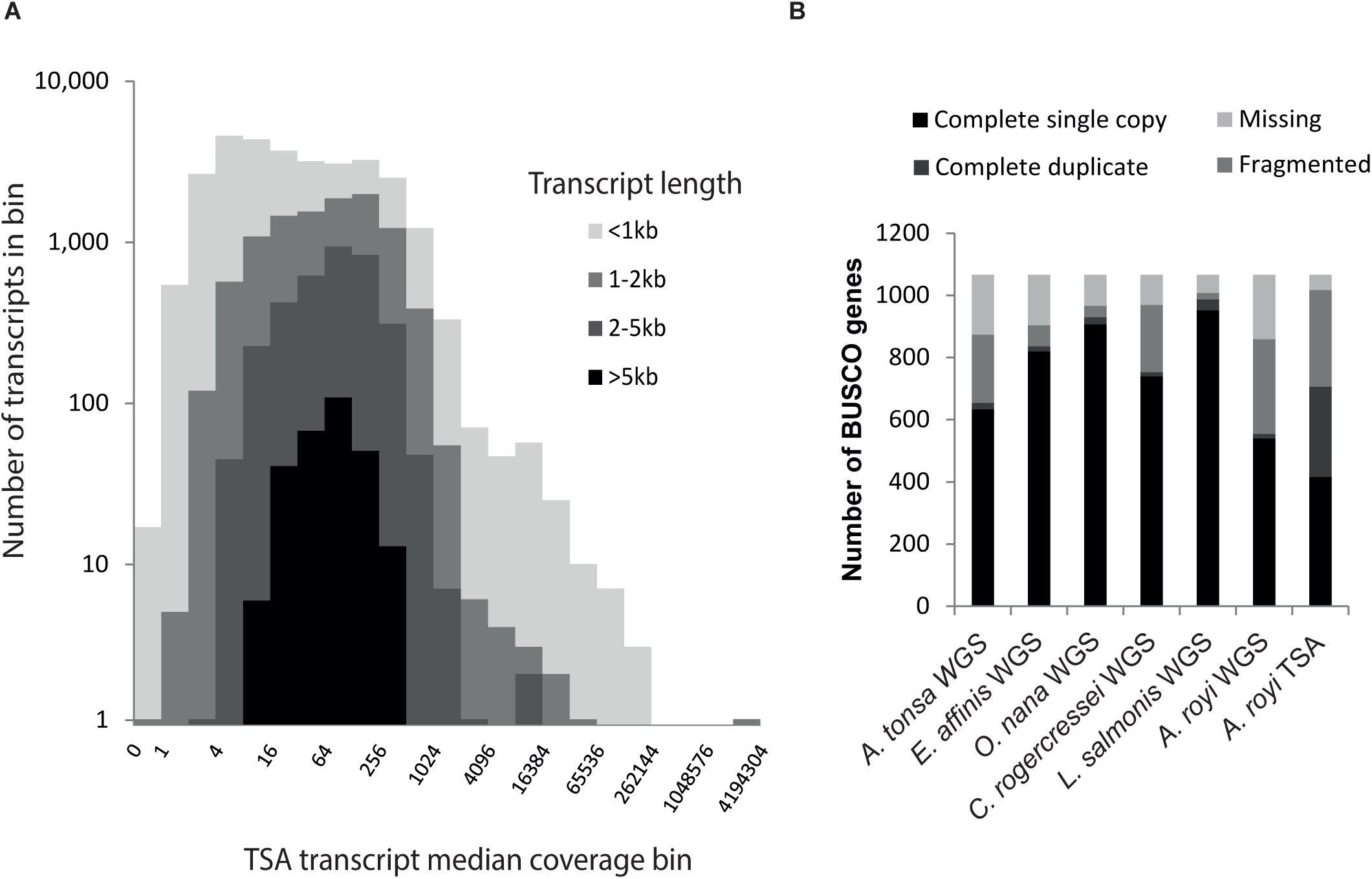
A, median coverage of TSA transcripts. While most transcripts have a median coverage of less than 200, several have median coverages of more than 100.000. The four length subset <1kb, 1-2kb, 2-5kb, and >5kb from the TSA are plotted, each with fewer transcripts, but each also with a similar distribution. Because the coverage of TSA transcripts is related to the expression of genes, large variations of the coverage of transcripts are both expected and observed. B, BUSCO scores for the available copepod WGS scaffolds (data from [13]) as well as the provided WGS and TSA assemblies of *A. royi*. For the *A. royi* WGS assembly, very few genes are duplicated (2%), while only just over 50% are complete and single copy. Another 29% are fragmented while 19% of genes are missing in the WGS assembly. For the TSA dataset, 66% of genes are complete, and another 29% fragmented while only 5% are missing. For the TSA data, the categories complete single copy and complete duplicate are not as important as for the WGS dataset. Importantly, in both the WGS and TSA datasets, only minimal fractions of the core BUSCO genes are not found at least in fragmented versions. This will allow using the information, though some analyses might be impaired.

In order to estimate the gene completeness of the WGS assembly, we used the BUSCO system of near-universal single-copy orthologous genes. We found 51 % complete and single copy genes, 1 % complete duplicated genes, 29 % fragmented and 19 % missing genes (Fig. 2B). These statistics are similar to some other copepod genome assemblies in the NCBI WGS database, and means that the large majority of conserved genes can be found in the assembly, though many are incomplete (Fig. 2B). The many fragmented genes could be explained by intron sizes up to 70 kb as recently reported in a crustacean [28]. For several practical applications, though, it is sufficient to have a gene fragment available to e.g. design primers for qPCR as long as it can be annotated unequivocally. An example of this from the presented TSA assembly can be found in a recent publication, where fatty acid desaturase genes were found in fragmented versions, reconstructed and which were found to be upregulated by starvation of polyunsaturated fatty acids in microalga feed [9]. For the TSA dataset, 706 BUSCO genes are complete (66 %), while another 311 BUSCO genes are fragmented (29 %) and 49 missing (5 %) (Fig. 2B). The complete BUSCO reports for the WGS and TSA assemblies are available in Supplementary Material 6 and 7. Almost all mitochondrial genes can be found on scaffold_16888 where only the ND4L gene and the small subunit of the ribosomal RNA gene are missing: the remaining 13 genes are all present as well as all 22 tRNAs, as determined by MITOS2 (data not shown) [29]. In order to phylogenetically place the presented *A. royi* WGS within the order Cyclopoida, we aligned the identified 18S rRNA gene sequence to the partial 18S rRNA gene sequences from a publication on the family level phylogeny of cyclopoid copepods [27]. The nucleotide sequence of the 18S rRNA gene can be found in Supplementary Material 8. The identified *A. royi* WGS 18S rRNA gene sequence shared 598nt out of 600nt with a database sequence from *Apocyclops royi* (acc.: HQ008747.1, data not shown). We then created a neighbor-joining tree and found that the *Apocyclops* sequences together with the sequence from our WGS data form a clade with high support (bootstrap values: 93% and 100%, Fig. 3). In general, readers are referred to [27] for a thorough phylogeny of cyclopoids as the branchings in Fig. 3 have little support. It does, however, thoroughly place the presented WGS assembly as *Apocyclops royi*.

**Fig 3.**
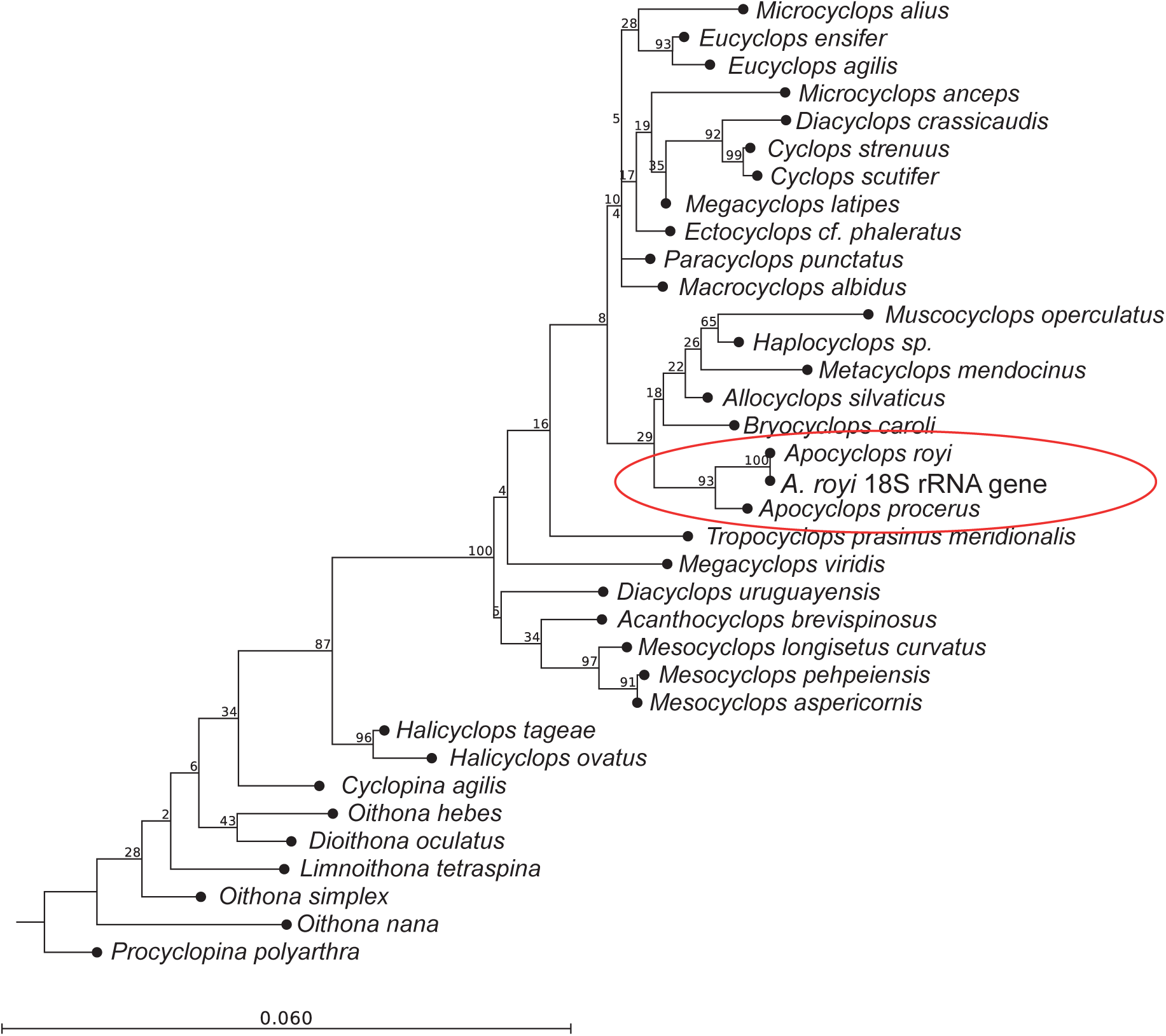
neighbor-joining tree with the cyclopoid 18S rRNA gene dataset from [27] and the identified 18S rRNA gene sequence from the presented WGS. While many branches have very low support, the sequences from the genus *Apocyclops* group together with high support (Bootstrap values of 93 % and 100 %). Further, the *A. royi* 18S rRNA gene sequence from the presented dataset is almost identical with the only sequence from database *A. royi*. This confirms the placement of the presented datasets as very closely related to the database *A. royi*.

In conclusion, we here present the WGS assembly (Acc. UYDY01) and an mRNA transcriptome assembly (Acc. GHAJ01) from the tropical cyclopoid copepod *Apocyclops royi*, along with the raw data used to produce them. We have shown that the provided datasets are sequenced to a sufficient depth, that any contamination in the raw reads has been removed from the WGS assembly, and that the phylogenetic placement within Cyclopoida matches our expectation for *Apocyclops royi*. Further, we have documented the completeness of core genes in both the TSA and WGS dataset and found 95% and 80% of core genes, though many in fragmented versions.

### Data availability statement

all raw data (Acc. ERR2811089, ERR2811715, ERR2811728-ERR2811734), the TSA assembly (Acc. GHAJ01), and the WGS assembly (Acc. UYDY01) are available in the ENA/NCBI system under project accession number PRJEB28764.

